# Inferring and simulating a gene regulatory network for the sympathoadrenal differentiation from single-cell transcriptomics in human

**DOI:** 10.1101/2025.03.21.644507

**Authors:** Olivier Gandrillon

**Affiliations:** Univ Lyon, ENS de Lyon, Univ Claude Bernard, CNRS UMR 5239, INSERM U1210, Laboratory of Biology and Modelling of the Cell, 46 allée d’Italie Site Jacques Monod, Lyon, F-69007, France; Inria, Villeurbanne, 69693, France

## Abstract

**Background:** Neuroblastoma is a malignant childhood cancer with significant inter- and intrapatient heterogeneity arising from the abnormal differentiation of neural crest cells into sympathetic neurons. The lack of actionable mutations limits therapeutic options, highlighting the need to better understand the molecular mechanisms that drive this differentiation. Although RNA velocity has provided some insights, modeling regulatory relationships is limited.

**Methods:** To address this, we applied our integrated gene regulatory network (GRNs) inference (CARDAMOM) and simulation (HARISSA) tools using a published single-cell RNAseq dataset from human sympathoadrenal differentiation.

**Results:** Our analysis identified a 97-gene GRN that drives the transition from Schwann cell precursors to chromaffin cells and sympathoblasts, highlighting dynamic interactions such as self-reinforcing loops and toggle switches. The simulation of that GRN was able to reproduce very satisfactorily the experimentally observed gene expression distributions.

**Conclusions:** Altogether, these findings demonstrate the utility of our GRN model framework for inferring GRN structure, even in the absence of a time-resolved dataset.

## Introduction

Neuroblastoma is a childhood cancer responsible for approximately 15 % of cancer-related deaths in children aged 0 to 4 years. These tumors typically develop near the adrenal glands and are characterized by significant inter- and intrapatient heterogeneity (1). Treatments are still limited, with few actionable mutations, and very few drugs have been clinically validated (2), although CAR-T cells targeting the GD2 antigen represent a promising prospect (3). Neuroblastoma is an archetypal example of a tumor arising through alterations in the differentiation process, more precisely during the differentiation of migratory neural crest cells into sympathetic neurons (4, 5, 6, 7). Therefore, there is an urgent need for a better understanding of the mechanisms driving normal development from neural crest and mesodermal lineages (8), to better understand the origin of different neuroblastoma subtypes, their internal heterogeneity, and interpatient variation in malignancy.

Initial studies were performed in mice (9). Although mouse models provide in-depth, experimentally supported insights into developmental processes, inconsistencies between mouse and human development can hinder our understanding of certain diseases. For example, transitions between cell fates during the differentiation of migratory neural crest cells into sympathetic neurons appear to occur in a different order in humans versus mice (10). Furthermore, neuroblastomas do not appear naturally in mice, and current genetic models do not recapitulate the full spectrum of natural properties of the disease (11). Therefore, it is critical to identify the human-specific aspects of development, which has led to a recent surge in studies of normal development at the single-cell level in humans (10; 12; 13). To go beyond a static description of the differentiation process, the authors of the aforementioned papers have resorted to the use of RNA velocity analysis to obtain a more dynamic view (10; 13). RNA velocity is driven by a kinetic model of RNA dynamics and works by distinguishing between unspliced and spliced mRNA counts and fitting gene-specific parameters based on an on-off deterministic transcription model. This allows the RNA velocity to predict the direction of future changes in spliced mRNA levels (14, 15, 16). However, although RNA velocity uses a time-dependent gene expression model, time has no explicit biological interpretation (17). Furthermore, each gene is modelled independently, and regulatory relationships are ignored (15).

We recently described a model for Gene Regulatory Networks (GRNs) that alleviates most of these drawbacks and provides detailed access to a fully mechanistic view of gene regulation during a differentiation process (18; 19). In this model, each gene is modelled as a piecewise deterministic Markov process (PDMP), describing bursts in the production of mRNA and the resulting synthesis of proteins (20; 21). Genes are coupled via an interaction function that describes how the proteomic field feeds back into burst frequency. The coupling of a GRN inference algorithm (CARDAMOM) with a simulation algorithm (HARISSA) allowed us to demonstrate the potential of this approach for inferring executable GRN models that can reproduce the observed experimental data.

In the present work, we describe the use of our integrated approach to infer the GRN driving sympathoadrenal differentiation in humans based on single-cell RNA-seq data (10), and to predict the effect of some gene perturbations. Our analysis identified a 97 genes-based GRN involved in the transition from Schwann cell precursors to chromaffin cells and sympathoblasts. Their study revealed dynamic gene interactions such as self-reinforcing loops and toggle switches.

## Material and methods

### Data preparation

The raw data is available at GEO GSE14782.

The processed adrenal.human.seurat.scrublet.rds dataset used in this study can be found at http://pklab.med.harvard.edu/artem/adrenal/data/Seurat/.

This represents single-cell transcriptomic analysis using 10x chromium on isolated individual cells from dissected adrenal glands with surrounding tissue from human embryos at 6, 8, 9, 11, 12, and 14. The initial matrix displays the expression of 25787 genes across 72571 cells. Only cells harboring a cell doublet probability (Scrublet score) < 0.2 were kept.

The cell types were annotated by the authors based upon gene expression pattern.

To elucidate the sympathoadrenal cell fate transitions in humans at a higher resolution, we focused on the 3,901 cells of SCP, chromaffin, and sympathetic fates, omitting annotated cell cycle genes. As advocated by the authors, we removed a small potentially contaminating population of cells expressing the STAR gene. This resulted in a final matrix displaying the expression level of 20807 genes in 3759 cells at six developmental time points. We verified that the UMAP generated from this dataset was in perfect agreement with Figure 2a in the original paper (see Figure 1a). UMAP was computed using the `Uwot` package (22). All the UMAPs were embedded within the experimental UMAP using a combination of `the ret_model` and `umap_transform` functions of the `uwot` package.

**Figure 1:**
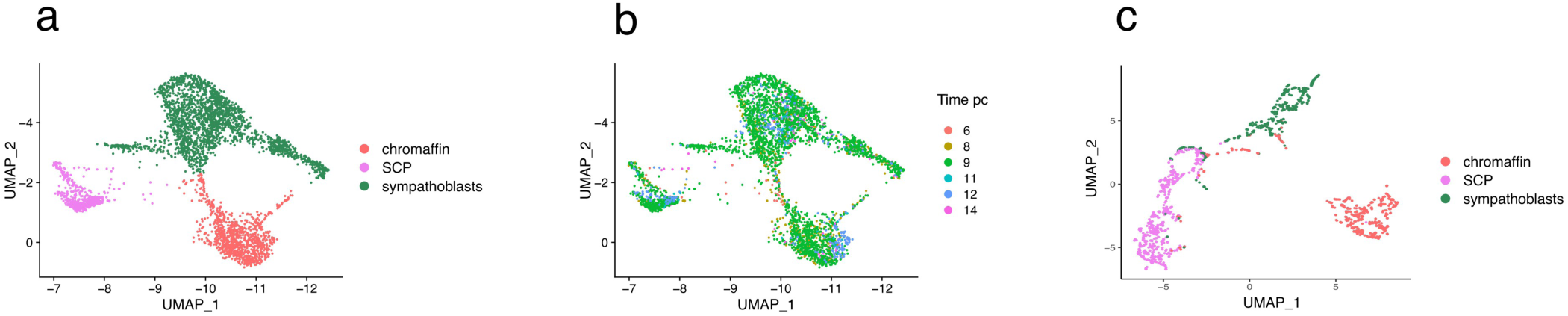
UMAP representation of the dataset. a and b: the initial 3759 cells x 20807 genes matrix. c: the final 1800 cell x 97 genes dataset after time sampling (see Figure 2). In a and c cells are color-coded according to their cell fate. In b, cells are color-coded according to their sampling time.

**Figure 2:**
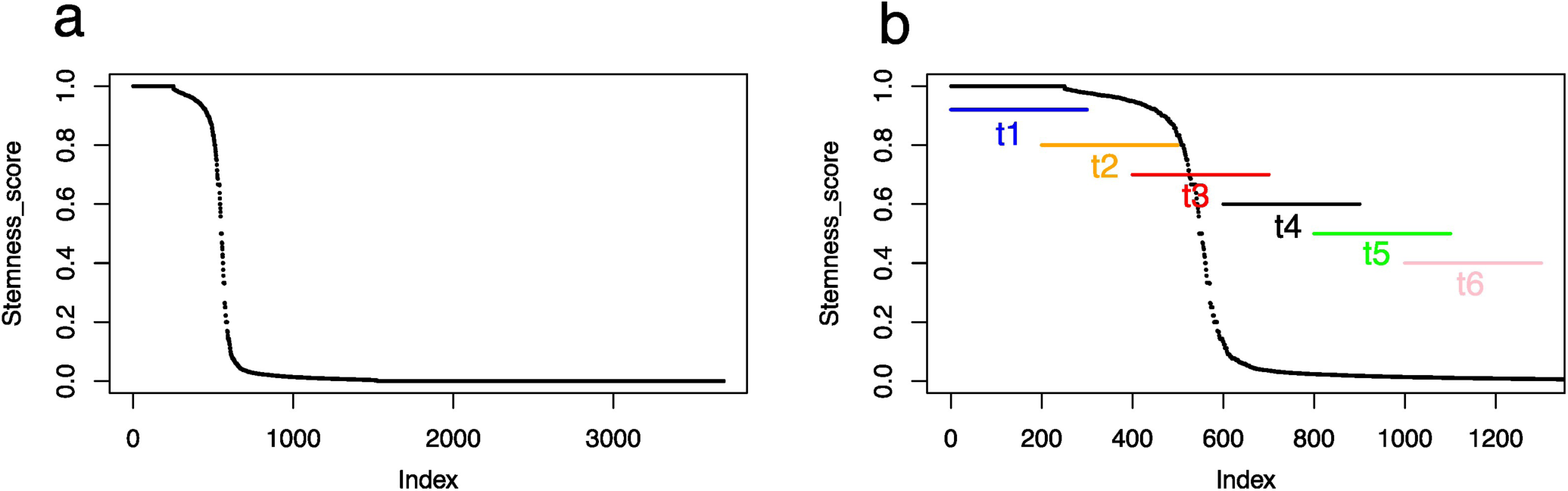
Cell ordering by their stemness score. a: all cells are displayed. b: Focus on the first 1300 cells, where the “process time” are indicated. The process time is defined along six time point with sliding windows of 300 cells overlapping by 100 cells.

### Stemness score

The stemness score was computed per cell as signature_SCP / signature_SCP + signature_sympatoblast + signature_chromaffin, by summing up the gene expression level for the following signature genes:

- SCPgenes: PLP1, FOXD3, FABP7, S100B, NGFR, ERBB3, MBP, MPZ, COL2A1, POSTN, MOXD1, and GAS7
- Sympatoblast genes: STMN2, HAND2, ELAVL4, STMN4, ISL1, PRPH, ELAVL2, and HMX1.
- chromaffin_genes: CHGA, CHGB, INSM1, PENK, PNMT and SLC35D3.

### GRN inference and representation

First, we cloned the original CARDAMOM git repository at https://github.com/ogandril/cardamom. All computations were performed at the IN2P3 computing center (https://cc.in2p3.fr). The scripts used for launching CARDAMOM on a distant server are available at https://github.com/ogandril/Launching_Cardamom. All interaction values were scaled by a factor of 10 and thresholded at a value of four to maintain only the most relevant interactions.

The resulting inter.csv interaction matrix was opened with R (23) and plotted using the `qgraph` package (24). A list of transcription factors was obtained from https://github.com/saezlab/CollecTRI (25). Violin plots were drawn using the `ggbeeswarm` package.

### GRN simulation

The simulation was performed using the CARDAMOM build-in version of HARISSA, a software for performing GRN simulations based on an underlying stochastic dynamical model driven by the transcriptional bursting phenomenon (26).

The dynamics of each gene is given by the following ordinary differential equations:

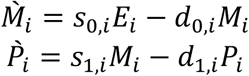

where M_i_ is the number of mRNA molecules for gene i, P_i_ is the number of proteins for gene I, E_i_ is the state of the promoter, s_0,i_ is the mRNA synthesis rate, d_0,i_ is the mRNA degradation rate, s_1,i_ is the protein synthesis rate, and d_1,i_ is the protein degradation rate.

E_i_ can randomly switch from 0 to 1 (activation) with a protein-dependent rate k_on,i_ (P) and from 1 to 0 (inactivation) with a constant rate k_off,i_.

We define the protein-dependent rate function k_on,i_ as in (26).

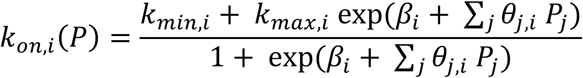

such that the promoter activation rate is between k_min,i_ and k_max,i_. The parameter β_i_ represents the basal activity of gene i, and the parameter 8_j;i_ encodes the value of the j → i interaction (see below for its dertemination).

Because we simulated a mechanistic model, one needs to provide HARISSA with half-lives for both mRNAs (d_0,i_) and proteins (d_1,i_). Experimentally determined values for mRNAs and proteins were extracted from (27) and (28), respectively. Genes that were not found in these datasets were attributed to the mean value of experimentally determined half-lives. Degradation rates were obtained using the following formula: degradation rate = ln(2)/half-life. The half-lives of proteins were limited by a parameter called the cell cycle, which was set to 20h.

Color code for a rapid estimation of the fit quality was as follows: given all Kantorovich distances for all genes at all times, we considered that 40 % of the values should be considered as correct (after a careful examination of the marginals) and therefore colored in green. The remaining 60 % was split into two, the first half colored in orange, and the last half colored red.

### Dynamical GRN representation

In CARDAMOM,we compute at each time point t_i_ with i ≥ 1 the value of each interaction, using

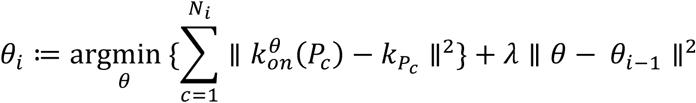

where Ni is the number of cells at time t_i_, P_c_ the amount of protein in cell c, *k_p_c__* the frequency mode and *θ*_0_ = 0.

So we do obtain at each time t_i_ a variation in the network

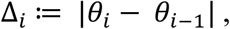

which quantifies how much each interaction estimate has been impacted by the cells measures at that time compared to all previous times. We can therefore, for any interaction between genes k and l, computes the times for which the largest impact was observed using

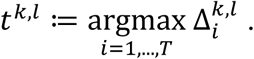

T being the latest time point available.

The dynamical interactions Θ_i_^k,l^ can therefore be defined as:

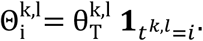

1 being the indicator function.

## Results

### 1. Organizing the cells

Our first task was to organize the cells according to their differentiation status, as our GRN inference is anchored in the time-dependent evolution of the differentiation process. Cells were collected from human embryos at 6, 8, 9, 11, 12, and 14 weeks post-conception. First, we assessed whether this cell collection time could be a relevant framework for generating time-reconstructed data. As shown in Figure 1b, this was not the case, since all cell types were present at all collection times. This is in line with the fact that in the human adrenal gland, the transition from Schwann cell precursor (*SCP*) to sympathoadrenal states persists for several weeks (10), which leads to the expectation that cells at all stages of differentiation should be present at all collection times.

Therefore, we were faced with a different situation from the one we used to benchmark CARDAMOM, where the cell sampling time was in accordance with the actual process time (18).

Consequently, we organized the cells according to their stemness scores (Figure 2a).

For this, we assumed a differentiation sequence starting with the SCP, giving rise to chromaffin cells and sympathoblasts through a bifurcation. Such a scheme is in line with the conclusions of the original paper from which we obtained dataset (10) and is shared by most (13, 12, 6, 29) but not all (4; 30) authors working on sympathoadrenal differentiation in humans. This led to the definition of the stemness score (see Materials and Methods) as the ratio of SCP-signature gene expression, evolving from 1 (only the SCP signature genes are expressed) to zero (none of the SCP signature genes are expressed). This allowed us to define the “process time” (Figure 2b) using an overlapping sliding window of 300 cells. This was defined only on the first 1300 cells since the stemness score was almost down to zero at the end of that sequence, and because we wanted to capture in detail the abrupt decrease in the stemness score. The use of an overlapping window is justified by the fact that differentiation should be considered a discontinuous nonlinear process (see e.g., 31), and therefore should allow cells to go back and forth in the gene expression space. This resulted in a 20807 genes x 1800 cells matrix.

### 2. Gene selection

We then explored the most relevant gene set to be used for the GRN inference. Because our inference and simulation scheme CARDAMOM is fundamentally rooted in exploiting the power of distributions (18; 32), we first computed a Kantorovitch distance (33) for all genes for all delta-times. Similarly, since most delta-entropic genes were shown to represent the most relevant genes for the differentiation process (34), we also computed delta-entropy for all genes for all delta-times.

The combination of these two lists (Figure 3) provided us with a list of 97 genes (Table 1) that were used for inferring the GRN.

**Figure 3:**
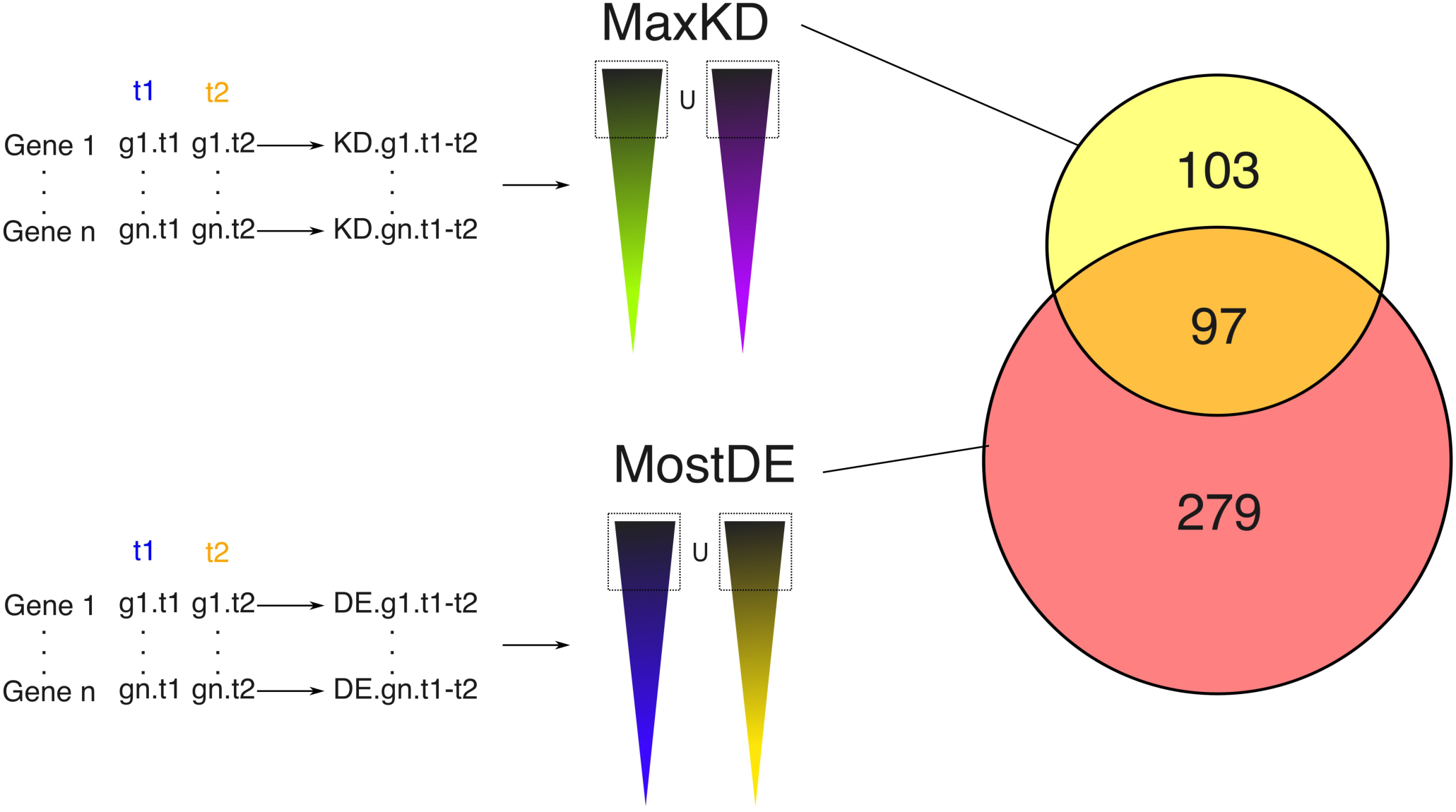
Gene selection procedure. We first computed a Kantorovitch distance for all genes between 2 consecutive time points. We then took the union of the 133 genes harboring the largest distance for each delta-time, which left us with a list of 200 genes (some genes might be among the most distant between two or more time points), labelled MaxKD. Very similarly, we computed an entropy value for all genes between 2 consecutive time points. We then took the union of the 50 genes harboring the largest entropy for each delta-time, which left us with a list of 376 genes labelled MostDE. The union of those two lists gave us a list of 97 genes (Table 1) that were used for inferring the GRN.

**Table 1:**
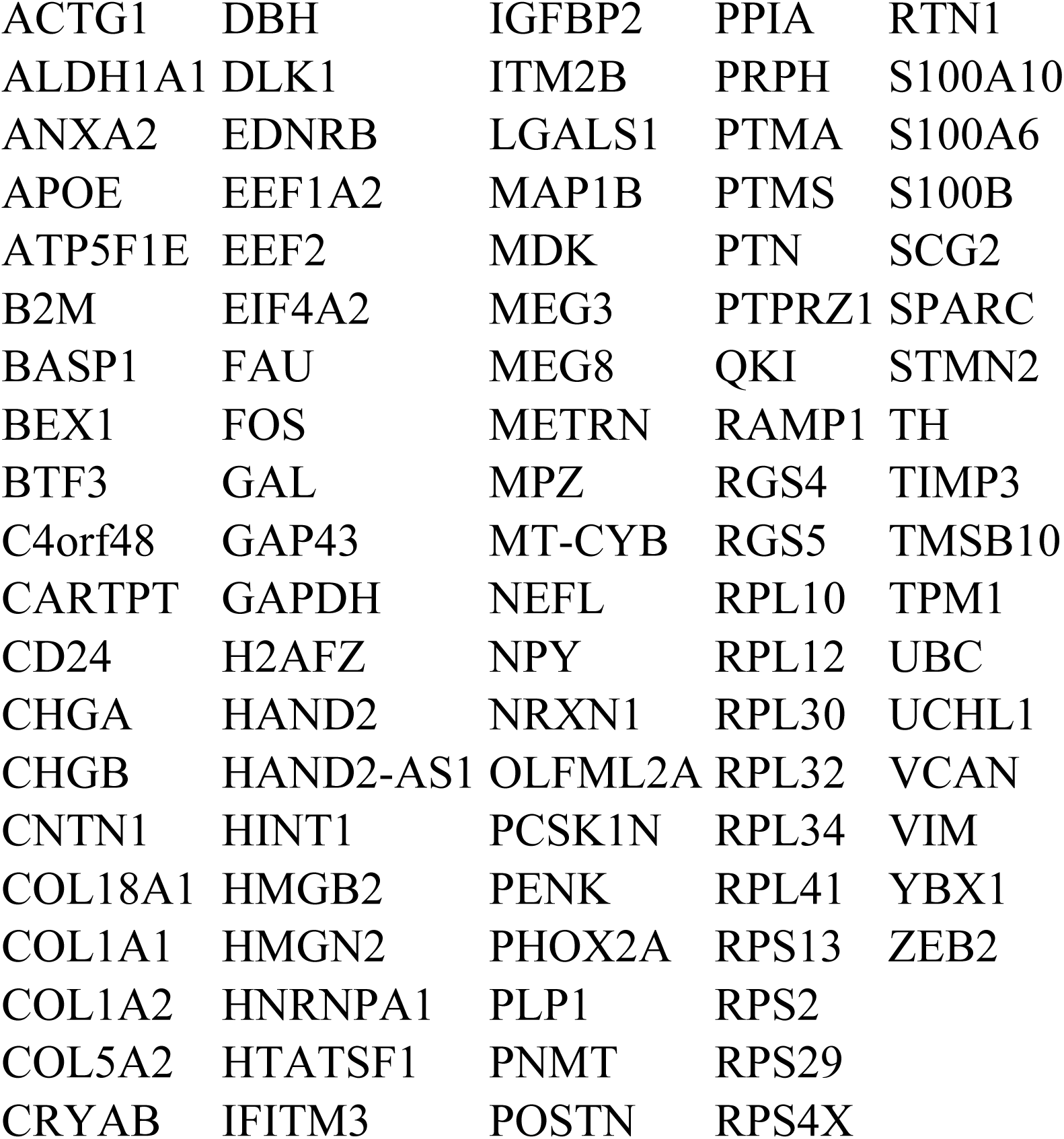
the 97 genes that were used for inferring the GRN.

This left us with a matrix displaying the expression levels of 97 genes in 1800 cells distributed over six time points. We verified that the information contained within this matrix was sufficient to correctly capture the differentiation process of interest, as assessed by UMAP representation (Figure 1c). It is important to note that these genes were chosen purely based on their expression patterns, irrespective of their putative biological functions (see Discussion).

### 3. GRN inference and analysis

We then applied the CARDAMOM algorithm to infer the GRN structure (Figure 4a), resulting in a relatively densely connected network.

**Figure 4:**
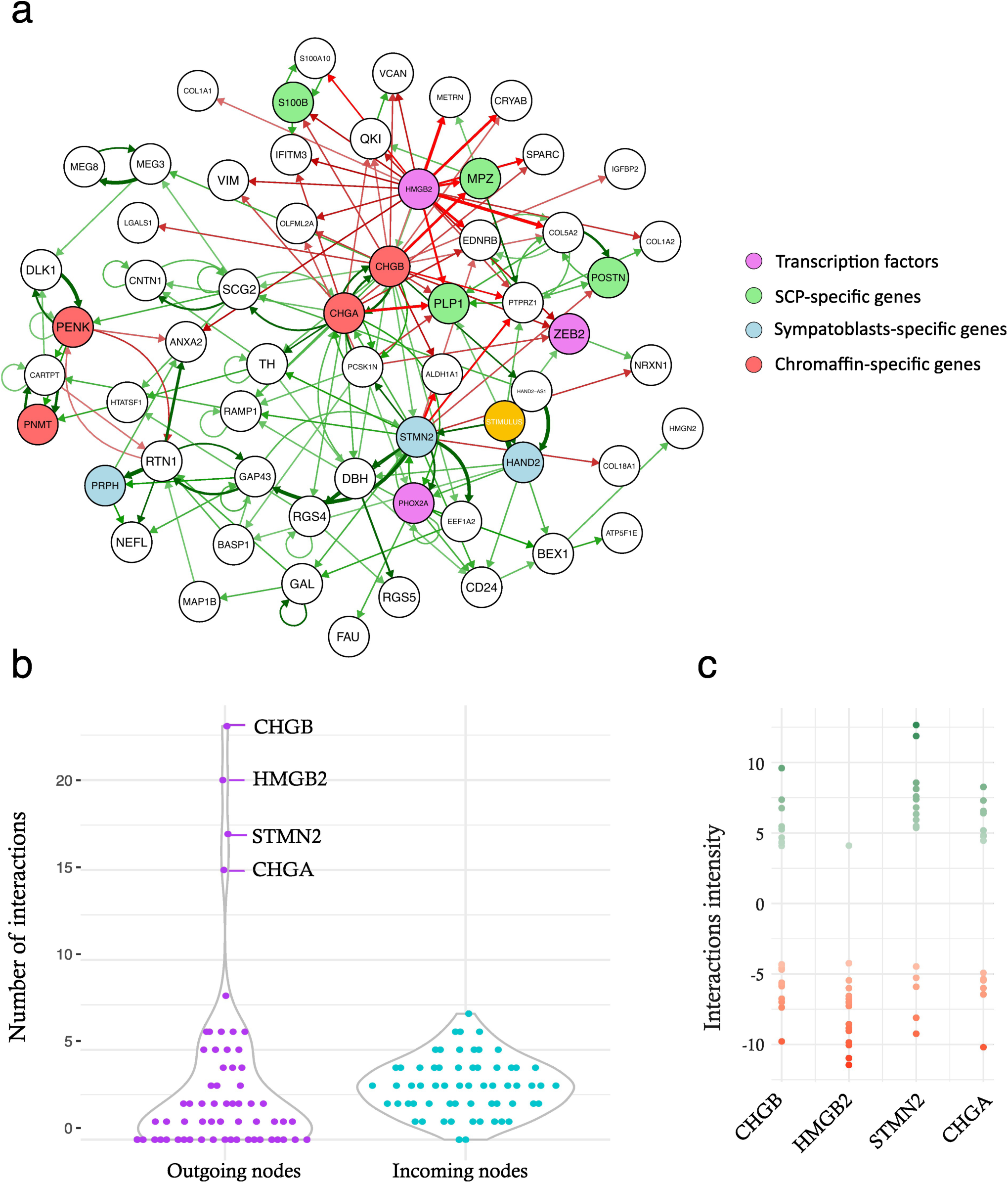
Representation of the inferred GRN structure. a: the full inferred GRN. Activations are displayed in green, inhibitions are displayed in red. The strength of the interaction is reflected by the arrow thickness. On the right is shown the color-code that has been applied to the GRN nodes. The stimulus is highlighted in yellow. b: violin plot representing the number of outgoing or incoming interactions per gene. c: the actual values of the outgoing interactions are shown for the four genes with the highest number of interactions.

First, we explored the network connectivity. We observed a striking difference between the distribution of the numbers of outgoing and incoming nodes (Figure 4b). The incoming nodes were distributed according to a normal (i.e., Gaussian) distribution centered around a mean of three, whereas the outgoing nodes displayed a very heavy long tail toward higher values, as well as a large number of null values (i.e., leaves, that is, genes only receiving and not sending any signals within the GRN). The long tail was mostly due to 4 genes (CHGA, CHGB, STMN2, and HMGB2). The intensity of all outgoing interactions for these four genes is shown in Figure 4c. HMGB2 displayed a specific pattern with quasi-exclusively inhibitory interactions, whereas the other three genes displayed a relatively similar number of positive and negative interactions. It should be noted that all but HMGB2 are part of the defining gene signatures (see Materials and Methods).

One key feature of our inference process is the possibility of decomposing the overall GRN into dynamical subparts, where each edge appears at the time point transition for which it was detected with the strongest intensity by the inference algorithm (Figure 5; 18, see Materials and Methods for a formal definition).

**Figure 5:**
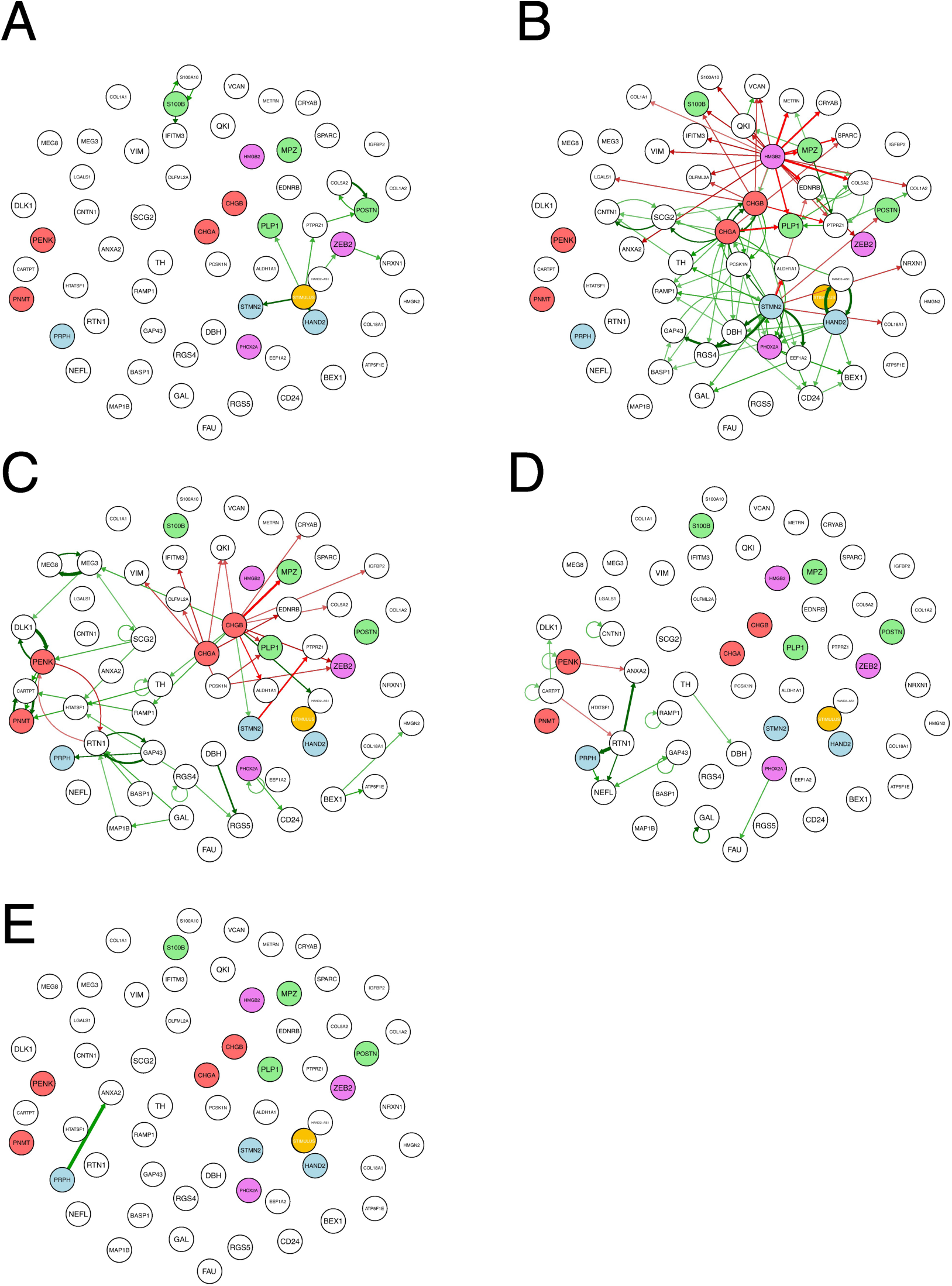
Representation of the time-dependent evolution of the inferred GRN structure. a: t1 → t2; b: t2 → t3; c: t3 → t4; d: t4 → t5; e: t5 → t6. For the time definition, see Figure 2. For the color-code applied to the GRN nodes, see Figure 4a. For the definition of the dynamical interactions displayed, see Material and Methods.

The dynamics start with a small number of genes being turned on by the stimulus (Figure 5a). In our previous studies, the stimulus was explicitly identified as an exogenous medium change/addition (35; 18). Here, it should be understood as a complex influence exerted by the cell environment, whether hormonal or through cell-cell interactions, which induces SCP cell differentiation. It is needed to “set the GRN in motion” and push the cells out of their quasi-steady state.

There was a much larger number of interactions important for the next time point (Figure 5b), including a very large set of genes being repressed by HMGB2. All known SCP-specific genes were repressed at that point, which was expected. A CHGA-CHGB self-reinforcing loop activates genes from the chromaffin lineage, while a few sympathoblast-specific genes (STMN2, HAND2) are activated.

At the next time point, an RTN1-PENK toggle switch was apparently supported by positive loops on both sides (Figure 5c). We can see a lot of genes being repressed by CHGA and CHGB genes. This includes the repression of MPZ-and PLP1 SCP-specific genes by CGHB, a chromaffin-specific gene.

The two latest time points showed a very small number of interactions (Figure 5d and 5e).

It is clear that the two time points where there is a large number of interactions correspond to the times at which one observes a steep variation in the stemness score (Figure 2). Altogether, this time decomposition offers a clear illustration of a signal propagating through the GRN in the form of waves of gene activation and repression (35).

Finally, we extracted from the full GRN a subset of the genes connected by specific dynamical motifs, that is, either cross-positive self-reinforcing loops or mutually repressive toggle switches (Figure 6). Among these, only one direct toggle switch can be observed, the one linking PENK to RTN1 genes. All other motifs were positive. One can see a group of genes (COL5A2, PTPRZ1, EDRNB, POSTN), including one SCP-specific gene (POSTN) which reinforce together and are repressed concomitantly by chromaffin specific genes (CHGA and CHGB) and a sympathoblast-specific gene (STMN2).

**Figure 6:**
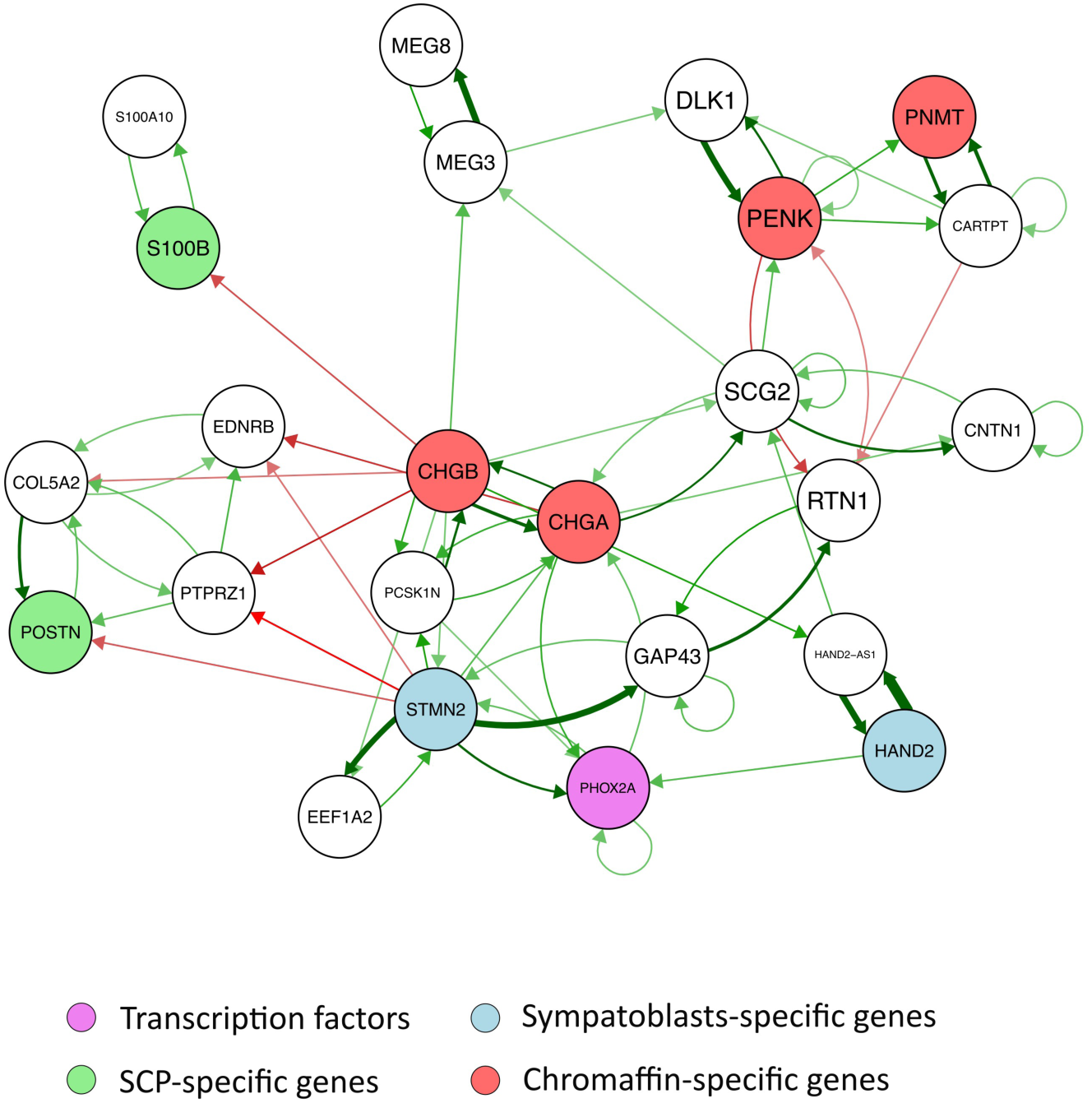
Representation of a subset of the GRN showing genes involved in specific dynamical motifs.

### 4. GRN simulation

One of the main advantages of our approach is that it produces executable GRN models (36), the time-dependent evolution of which can be simulated, and the resulting simulated dataset can be compared with the experimentally observed dataset.

There are many ways in which the results of a simulation can be assessed. Using a UMAP representation allows for quick evaluation of the quality of the fit. This was first used to calibrate the process times t1– t5 in real time (Figure 7).

**Figure 7:**
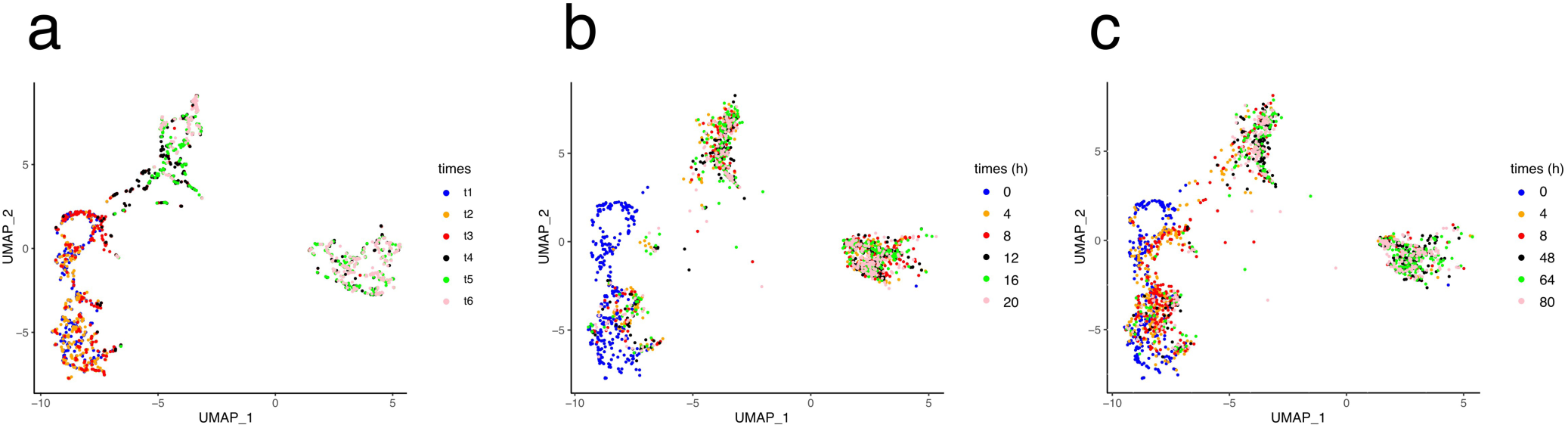
Adjusting the process time to real time values. In a, the UMAP is computed on the experimental dataset and color coded according to the process times (see Figure 2). b and c: the UMAP is computed on the simulated dataset and color coded according to the model time, in hours.

We first tested an evenly spaced time frame, where each time point was set 4 h apart (Figure 7b). All the cells were instantly projected from their t1 position to the final position of the process. Therefore, we attempted different adjustments and obtained the best visual fit using a time sequence extending up to 80 h (Figure 7c), which was used for further simulations.

We then assessed the extent to which the simulation captured the time-dependent evolution of a few genes whose expression was characteristic of the three cell types we were trying to reproduce (Figure 8).

**Figure 8:**
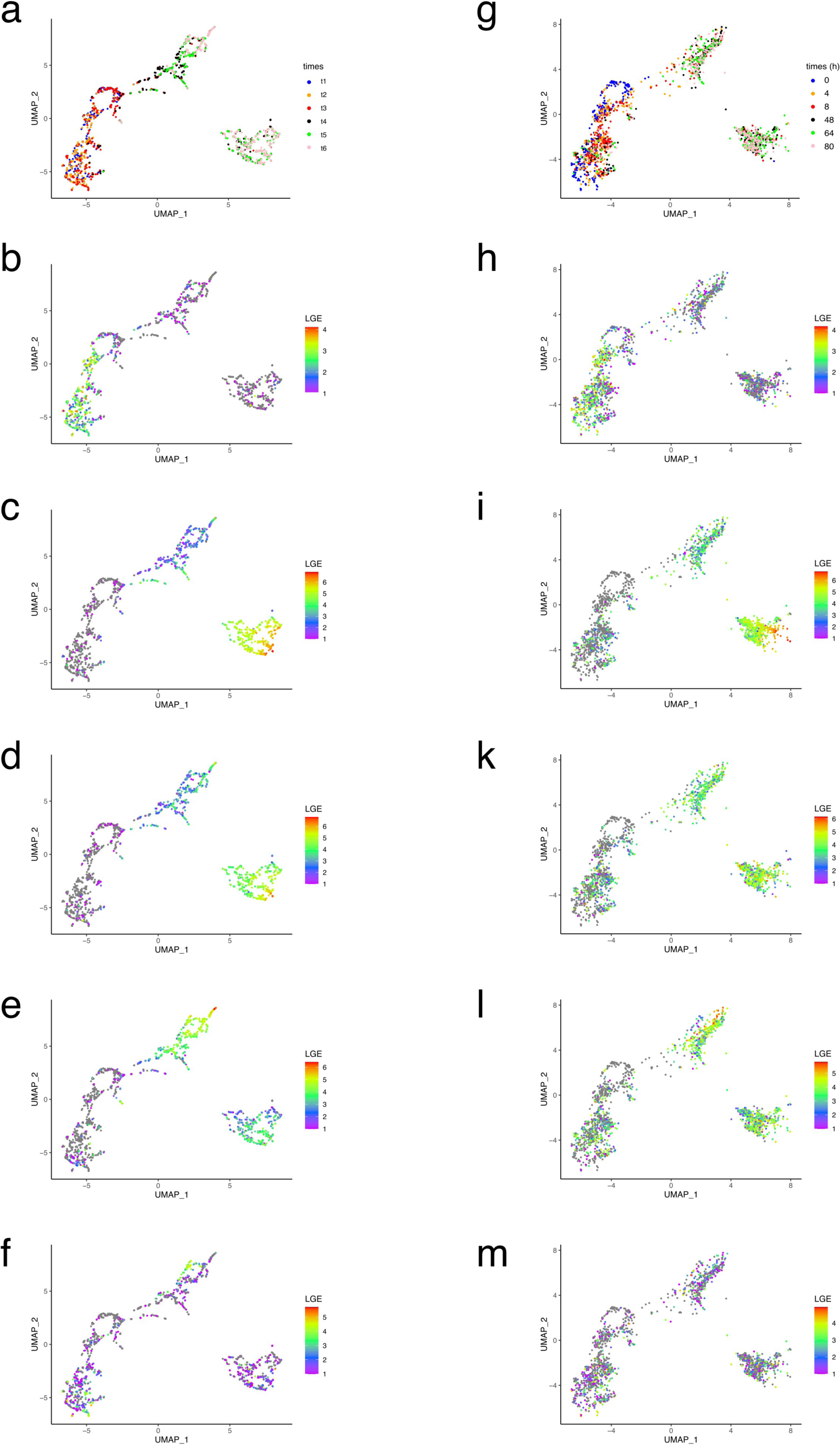
UMAP representation of the dataset used for GRN inference (a-f) and of the dataset obtained through GRN simulation (g-m). The cells are colored coded according to: (i) time (a and g); (ii) ZEB2 expression level (b and h); (iii) CHGA expression level (c and i); (iv): CHGB expression level (d and k); (v): STMN2 expression level (e and l) and (vi): HMGB2 expression level (f and m). All gene expression are log-transformed (LGE= ln(x+1)).

Overall, the gene expression pattern, when observed through its UMAP projection, was quite well reproduced by our GRN simulation. Most notably, chromaffin/sympathoblast bifurcation was quite well captured by our model.

Another way to assess the quality of the fit is to investigate the distance between the distribution of gene expression values of the experimentally observed mRNA distribution and the simulated distribution. As shown in Figure 9, there was a close proximity between the two distributions, as assessed by a low Kantorovich distance for the four genes displayed. Altogether, although some refinement could be performed (see discussion), we considered the fit quality of the experimental dataset to be quite satisfactory.

**Figure 9:**
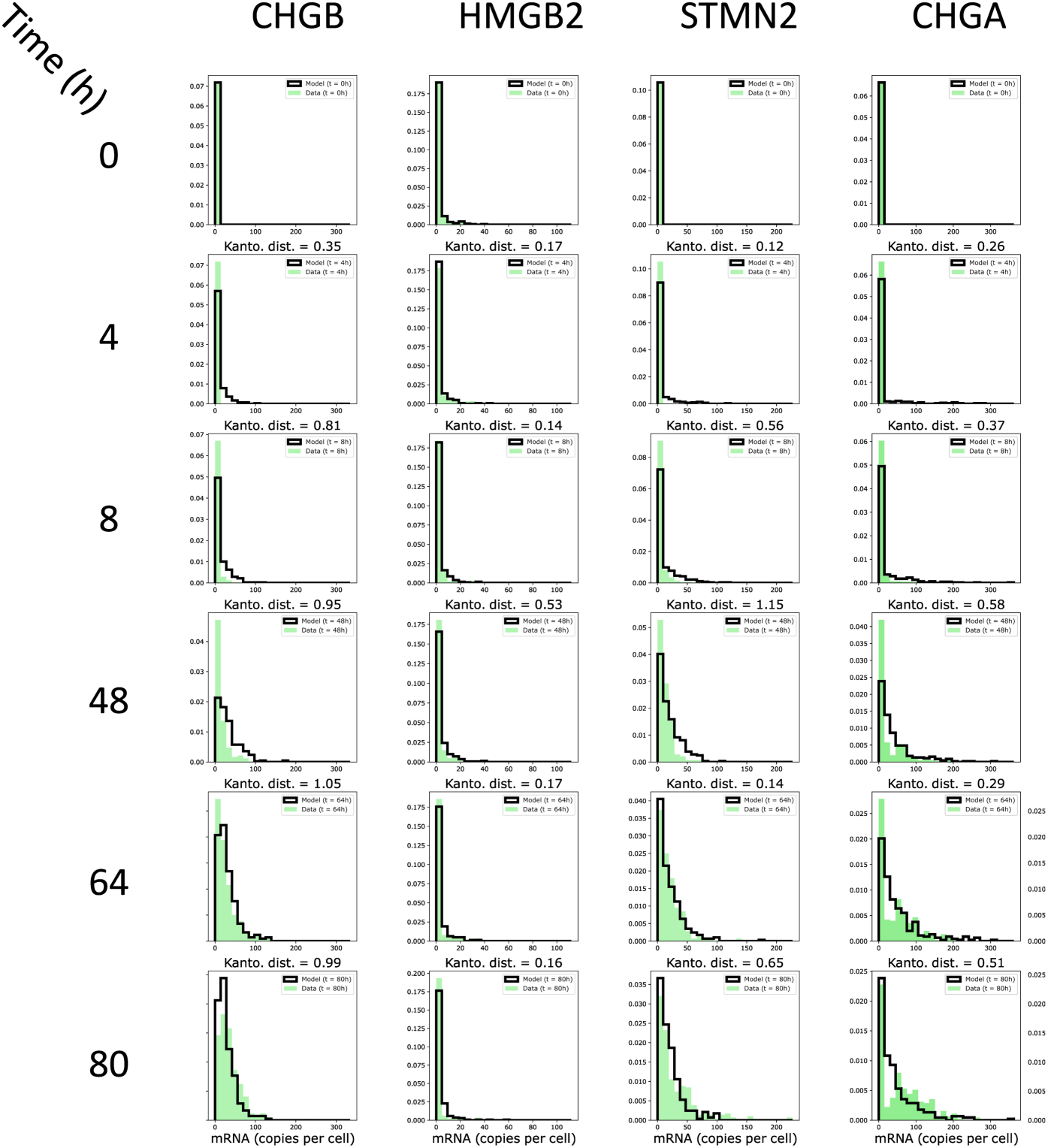
Fit quality for each gene at each time point as assessed by the Kantorovich distance (Kanto.dist) between the experimentally observed mRNA distribution and the simulated distribution. The marginals computed by the model are shown by bold-delimited bars, the experimenatlly-observed margnials are shown as plain bars in green.

We finally analyzed the run-to-run variability of the model, given the intrinsically stochastic behavior of our gene expression model (see “GRN simulation” upper). We therefore performed 10 runs of the GRN without specifying the seed, and analyzed the result by taking advantage of the clear separation of the three cell types in the 2D UMAP space (Figure 1c). This allowed us to count the number of cells belonging to each cell type following the simulation (Figure 10). We observed a very reproducible behavior of our model with only marginal differences between the number of cells for each cell type between different simulations.

**Figure 10:**
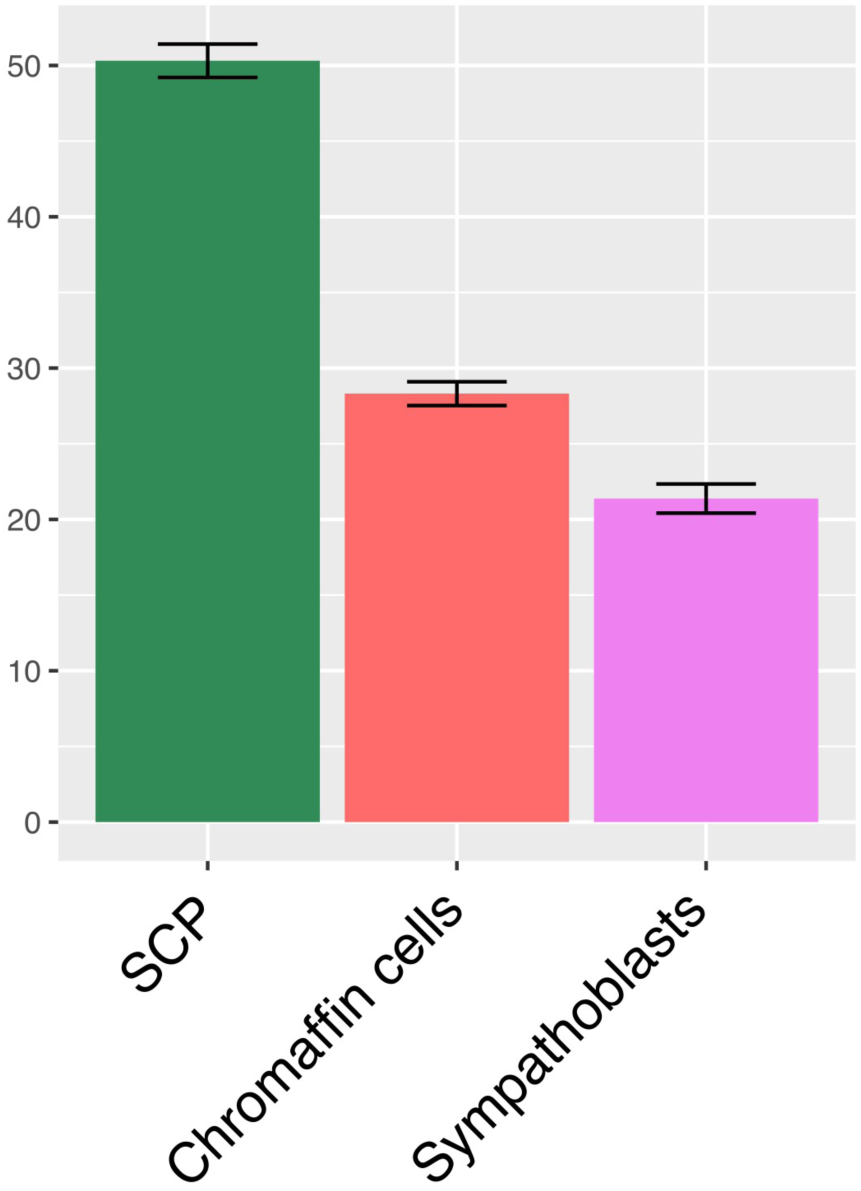
The UMAP space was divided in three regions, representing SCP, sympathoblasts and chromaffin cells (see Figure 1c). Shown the mean +/- SD of the percent of cells belonging to each cell category, as assessed by their 2D UMAP position, in 10 independent runs of the model.

## Discussion

One of the difficulties we had to solve in the present work is that the sampling time is not equal to the process time because all differentiation steps appear at all collected time points. As previously mentioned, “our approach can be applied to any biological process in which time-stamped single-cell transcriptomic data are obtained after applying a given stimulus. When such time-stamped snapshots are not available [which is the case here], the algorithm could, in principle, take as input time-reconstructed data (i.e., artificially ordered snapshots). In this case, the quality of the inference strictly depends on the effectiveness of the time reconstruction algorithm.” (18).

Therefore, we assumed a differentiation sequence starting with the Schwann cell precursor (SCP), giving rise to chromaffin cells and sympathoblasts through a bifurcation. This allowed the organization of cells along a stemness score aimed at capturing the unfolding of the differentiation process. This differentiation sequence was demonstrated in the study in which we obtained our dataset (10). Such a sequence is shared by some authors (13, 12, 6) but not all (4; 30). In the last case, the authors assumed a differentiation sequence starting with sympathoblasts and giving rise to chromaffin cells and SCP through a bifurcation. The reason for such discrepancies is unclear and sheds a crude light on the complexity of ascertaining a differentiation sequence *in vivo* in humans. One must stress that our method relies on a given definition of the differentiation sequence and cannot be used to untangle different hypotheses.

Assuming this order in the differentiation sequence, we proposed a framework through which cells could be organized, genes could be selected, and a 97 genes-based GRN could be inferred.

When compared to the GRNs previously inferred on different differentiation sequences (35; 18), the structure of the adrenal differentiation GRN displays both similarities and differences. Among these differences, one should note that the depth of the GRN (i.e., the largest number of nodes connected in a row) is slightly larger (6) than that for the erythroid differentiation sequence (depth of 3) or for the RA-induced ES cell differentiation (depth of 4). This is not imposed by our inference algorithm and can potentially infer a GRN of any length. In any case, this depth is limited by the duration of the differentiation sequence, which is in the same range (80 h) as that for the erythroid (72 h) or ES (96 h) differentiation sequences. If the network depth is too large, the signal is too damped and delayed to reproduce the experimental data accurately. Another difference was the role played by the stimulus; sympathoblast differentiation displayed a relatively modest effect (four genes affected directly by the stimulus), whereas it had a much more pronounced effect on ES (14 genes affected) and erythroid (29 genes affected) differentiation sequences.

A very clear similarity in all GRN behaviors is that there is a very clear wave-like pattern in the way the signal propagates through the network. In the case of sympathoblast differentiation, the more dense interaction times are seen at the t2-t3 and t3-t4 transitions points (Figure 5), in line with the very strong and abrupt change in stemness observed at those time points (Figure 2b).

The analysis of the subset of genes connected by specific dynamical motifs showed the existence of a group of genes, including two signature SCP-specific genes (POSTN and S100B) that reinforce together and are repressed by chromaffin-specific genes and a sympathoblast-specific gene. This is reminiscent of the “team” concept: a team is a group of genes that activates member genes belonging to the same team while inhibiting genes of other teams directly and/or indirectly. It has been shown to be a network design principle that can drive cell fate canalization in diverse decision-making processes (37) in a robust manner (38). In our case, this could have been involved in the maintenance of an undifferentiated SCP phenotype, which would have to be turned down to allow differentiation to proceed.

This study aimed to infer and simulate a GRN operating during normal sympathoadrenal differentiation in humans. The main incentive behind the study of this specific differentiation sequence is to provide insights from human developmental studies relevant to cases of neural crest-related pathologies and cancers. Therefore, our work is potentially an appropriate step in the understanding the developmental origin of neuroblastoma. It has been suggested that some immature sympathetic cells or their immediate progenitors may be of tumor origin (5). Nevertheless, the current ambiguity regarding what should be considered in the actual differentiation sequence (13, 12, 6, 4, 30) complicates this debate. It is clear that the generation of neuroblastoma cells cannot be resumed as a simple differentiation blockade in an immature SCP state.

However, there are clear limitations to this approach. Although we describe an informed gene selection procedure, it is dependent upon the use of a given arbitrary threshold for the number of genes selected. The use of a double selection procedure (using both Kantorovich distances and entropy) might alleviate this effect, but we cannot rule out that selecting more genes would have led to the incorporation of biologically relevant genes. We would also like to emphasize that contrary to most, if not all, GRN inference algorithms our gene selection procedure is completely blind to the function of the selected genes and relies purely on the time-dependent evolution of their mRNA distributions. This implies that many genes selected are not transcription factors, a very clear difference from most GRN inference, where regulatory proteins within a GRN are restricted to transcription factors (TF), as in (39; 40; 41). Possible indirect interactions were completely ignored. A trivial example is the gene encoding a protein that induces the nuclear translocation of a constitutive TF. In this case, the regulator gene indirectly regulates TF target genes, and its effect is crucial for understanding GRN behavior. In our case, assuming a time-scale separation, we reasoned that only the genes that displayed a change in their expression pattern (i.e., their mRNA distribution at the single-cell level) within the time frame of the experimental sampling could be captured and used for GRN inference, based on single-cell transcriptomics. This means that most of the interactions that our GRN display are probably indirect and are molecularly mediated by changes that act at scales that are not captured by the sampling process (e.g., very fast protein phosphorylation). Therefore, edges in our GRN should be interpreted with caution, and elucidating their molecular nature would require dedicated efforts using different techniques and sampling strategies.

One of the main difficulties in correctly reproducing experimental data using a mechanistic model is that critical values regarding the dynamical behavior of the model, that is both mRNAs and proteins half-life, have to be feeded to the model, based either on literature or through a rough estimate. Therefore an algorithm that would infer the half-life values while inferring the GRN structure would represent an invaluable step forward compared to the actual CARDAMOM algorithm.

In summary, we have proposed an approach for inferring and simulating GRN based on single-cell transcriptomic data generated to sample an *in vivo* differentiation process. We showed that the absence of a time-stamped process in a dataset can be overcome by *ad hoc* measures. There is little doubt that, given the flood of such a dataset in the literature, the use of our approach could lead to a better understanding of the mechanisms operating during cell decision-making.

## Ethics approval and consent to participate

Not applicable

## Consent for publication

OG approved the publication

## Availability of data and material

CARDAMOM is available at https://github.com/ogandril/cardamom. The scripts used for launching CARDAMOM on a distant server are available at https://github.com/ogandril/Launching_Cardamom.

No data associated with this article

## Authors’ contributions

OG analyzed the dataset, generated and analyzed the GRN, performed the in silico simulations, and wrote the manuscript.

## Acknowledgements

I am grateful to Elias Ventre, Ulysse Herbach, and Matteo Bouvier for their continuous support and help in setting up and improving the CARDAMOM/HARISSA framework.

I thank Elias Ventre and Sandrine Gonin-Giraud for their critical reading of the manuscript.

I thank the computational center of IN2P3 (Villeurbanne/France), where the computations were performed. I thank the BioSyL Federation and LabEx Ecofect (ANR-11-LABX-0048) of the University of Lyon for inspiring scientific events.

This work was not financed by the Association Pour la Recherche Contre le Cancer (ARC). A preliminary version of this work is available at : https://www.biorxiv.org/content/10.1101/2025.03.21.644507v2.

## Software Availability

Source code available from: https://github.com/ogandril/cardamom and https://github.com/ogandril/Launching_Cardamom

Archived software available from: https://doi.org/10.5281/zenodo.15389612 and https://doi.org/10.5281/zenodo.15389635

License: CeCILL Free Software License Agreement v2.1

